## Supplemental Figures for "Inversions Can Accumulate Balanced Sexual Antagonism: Evidence from Simulations and Drosophila Experiments"

### Supplemental Figure 1

Single locus simulations under differing models exhibit shifts in the parameter combinations that produce balancing selection. Each pixel in the heat-map plots represents 100 simulations of a population with a single polymorphic mutation of a given survival probability and display value that began at 0.5 frequency. Plots show either the average frequency of the mutation across simulations at 20*N* generations (left panels; green indicates intermediate balanced frequencies), or the proportion of simulations retaining polymorphism at 20*N* generations (right panels; dark blue indicates all or nearly all simulations retaining polymorphism). Plots are separated into columns based on whether the simulations assign survival costs to both sexes or to either sex individually. Plots are separated into rows by the number of males ***m*** encountered by each female during mate choice. Simulations have a population size of 1,000 individuals simulated for 20,000 generations, with further details as indicated in the Materials and Methods. Discussed in relation to Figure 3 in the main text.

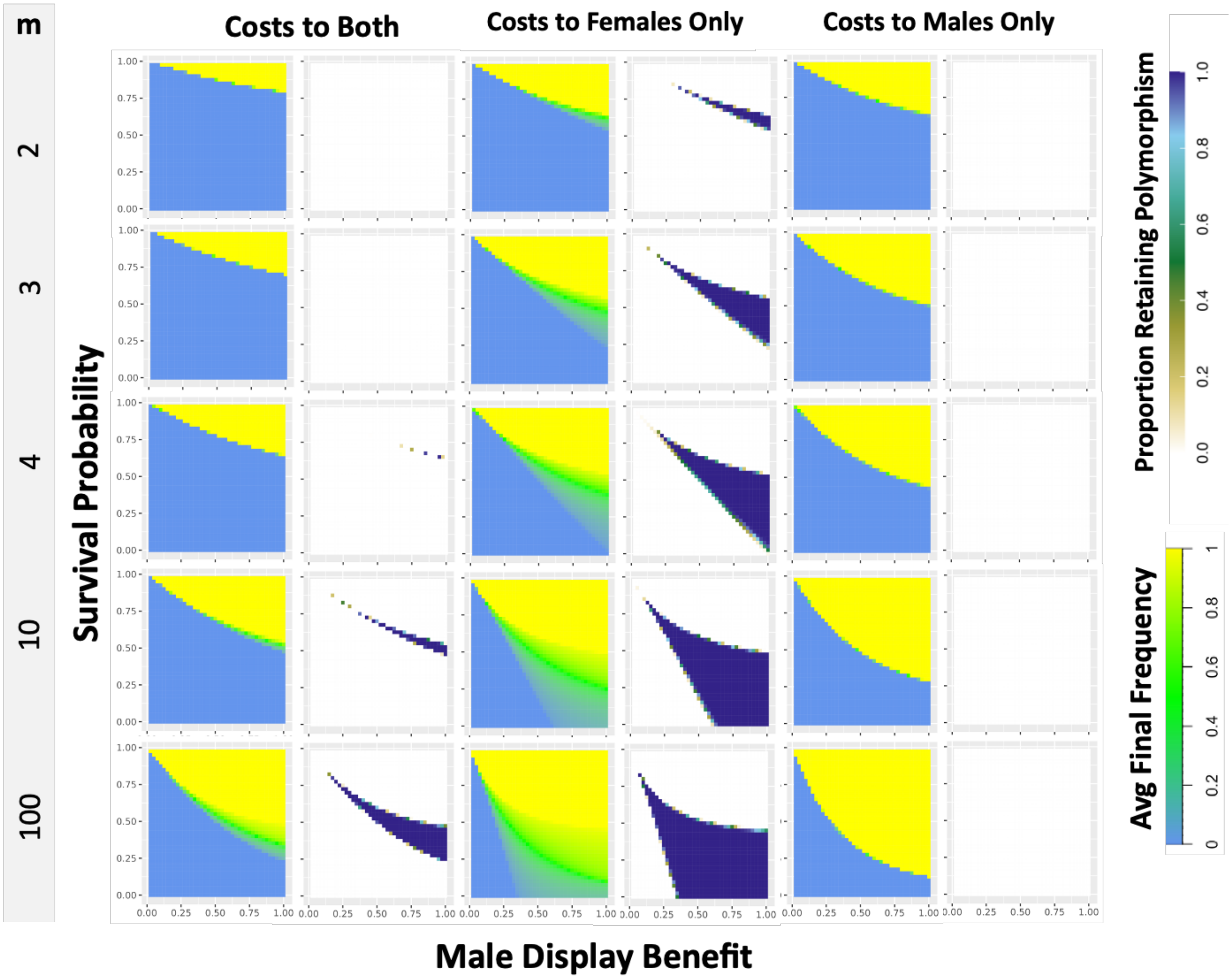
m
2
3
4
10
100
Costs to Both
Costs to Females Only
Costs to Males Only
Survival Probability
Male Display Benefit
1.00
0.75
0.50
0.25
0.00
0.00 0.25 0.50 0.75 1.00
Proportion Retaining Polymorphism
0.0
0.2
0.4
0.6
0.8
1.0
Avg Final Frequency
0
0.2
0.4
0.6
0.8
1

### Supplemental Figure 2

After 20N generations of simulation, populations approach an equilibrium in which the haplotypes with both display-favoring or both survival-favoring variants are most fit, and selection removes recombinant haplotypes, maintaining high linkage. (A) The number of descendant haplotypes per haplotype for each haplotype class averaged across all 1000 simulations, plotted for both sexes and within females and males separately. From the reproductive values across both sexes (1.0038, 0.9793, 0.9926, 1.0047) haplotypes with both display-favoring or survival-favoring variants are favored over haplotypes conveying intermediate trait values. (B) The average frequency of each haplotype class in its population across simulations. Haplotypes with intermediate trait values are maintained at low frequency under a balance of recombination and selection (0.3433, 0.0728, 0.2093, 0.3746), generating considerable linkage disequilibrium ($r^2$=0.2139). These simulations reflect one of the scenarios depicted in Figure 4A, in which the larger and smaller effect alleles were positioned at 0.225M and 0.250M respectively, and no inversion was present.

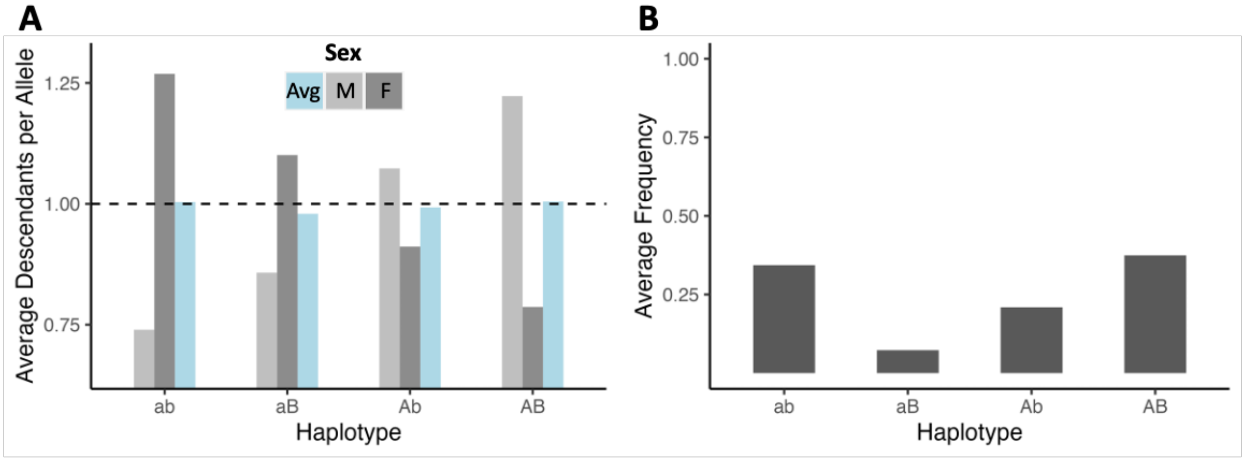

### Supplemental Figure 3

A histogram of the observed inversion frequencies in simulations with mutational generation of inversions with random position and length. In these simulations, only one inversion was allowed per haplotype, to enable easier accounting of the average effects of variants associated with an arrangement. Without antagonistic mutations, or other fitness effects, the distribution of inversion frequencies across all simulation replicates reflects the distribution expected for a neutral variant (left panel), although this figure represents the frequency of inversions across all simulations instead of a single population because only a couple inversions are expected in each simulation run. With antagonistic mutations, inversions tend to cluster around frequencies near 0.25 and 0.75 (right panel), due to parental transmission dynamics resembling sex chromosomes. Discussed in relation to Figure 6 in the main text.

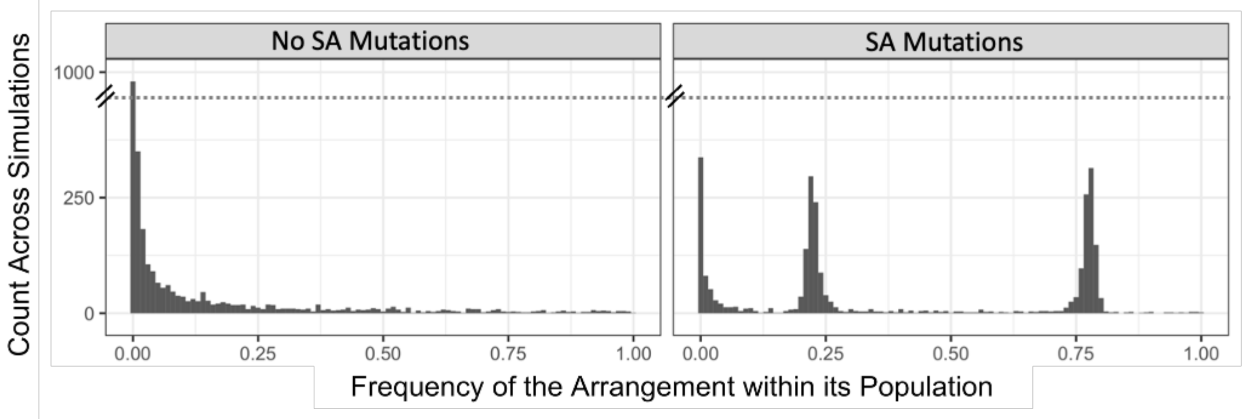

### Supplemental Figure 4

A plot of the observed inversion frequencies in early-eclosing and late-eclosing adult offspring cohorts, compared to the embryo frequencies. Inversion frequency changes were largely parallel between these cohorts, and no significant effects of eclosion time were identified across maternal line cross replicates. Discussed in relation to Figure 9 in the main text.

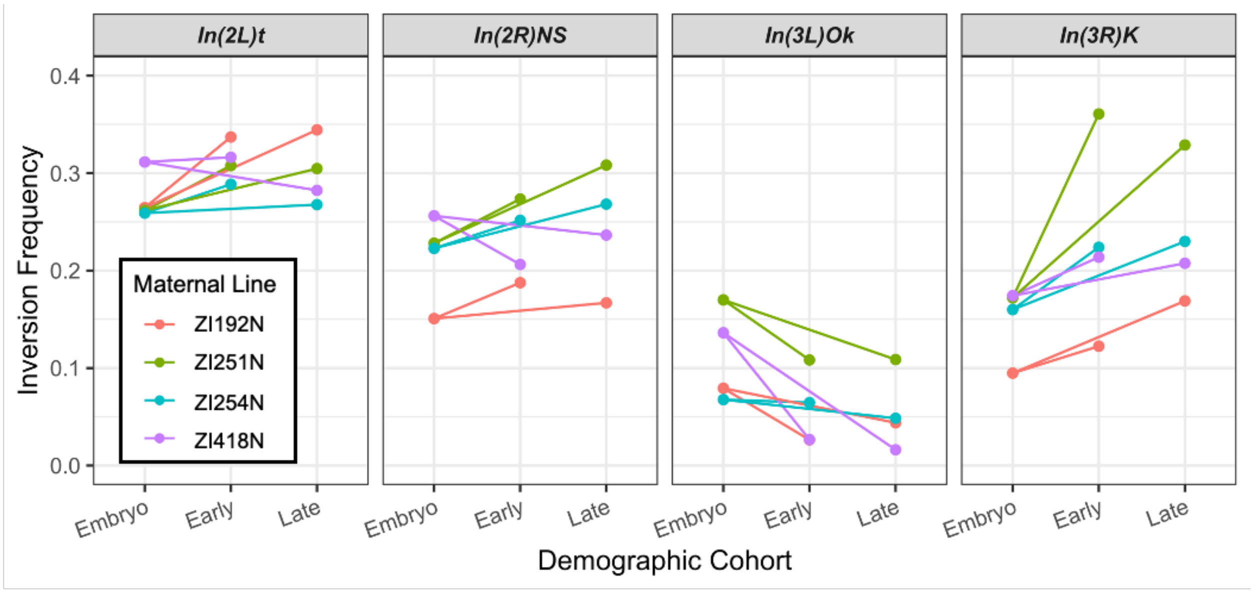
